# Fragile X mental retardation protein is a size-dependent translational activator

**DOI:** 10.1101/257204

**Authors:** Ethan J. Greenblatt, Allan C. Spradling

**Affiliations:** Howard Hughes Medical Institute Research Laboratories, Department of Embryology, Carnegie Institution for Science, 3520 San Martin Dr., Baltimore, MD 21218

**Keywords:** FMR1, mental retardation, autism, ovarian follicle, translational regulation

## Abstract

FMR1 enhances translation of large neural/oocyte proteins

Mutations in the highly conserved *Fragile X mental retardation gene (Fmr1)* cause the most common inherited human intellectual disability/autism spectrum disorder. *Fmr1* is also needed for ovarian follicle development, and lesions are the largest genetic cause of premature ovarian failure (POF). FMR1 associates with ribosomes and is thought to repress translation, but identifying functional targets has been difficult. We analyzed FMR1’s role in quiescent *Drosophila* oocytes stored prior to ovulation, cells that depend entirely on translation of stored mRNA. Ribosome profiling revealed that in quiescent oocytes FMR1 stimulates the translation of large proteins, including at least twelve proteins whose human homologs are associated with dominant intellectual disability disorders, and 25 others associated with neural dysfunction. Knockdown of Fmr1 in unstored oocytes did not affect embryo development, but more than 50% of embryos derived from stored oocytes lacking FMR1 developed severe neural defects. Fmr1’s previously unappreciated role promoting the translation of large proteins from stored mRNAs in oocytes and neurons may underlie POF as well as multiple aspects of neural dysfunction.

---

FMR1 is a polysome-associated RNA-binding protein (*1*-*5*) required for the ovary, testis and nervous system to develop and function normally in humans, mice and *Drosophila* (*6*-*12*). All three tissues rely heavily on the translational control of stored mRNAs associated with ribonucleoprotein particles (RNPs) containing FMR1 and conserved proteins such as Staufen, Pumilio and Ddx6/Me31B (*13*-*16*). FMR1-rich RNPs are found near synapses in neurons (*13*, *17*, *18*) and throughout mature *Drosophila* and human oocytes (*19*, *20*), suggesting that FMR1 specifically functions in utilizing stored mRNAs. However, the localized and dynamic nature of translational control at neuronal synapses, and the challenge of obtaining material that is highly enriched in FMR1-containing RNPs may have contributed to the difficulty in defining FMR1 target genes (*21*-*23*). We reasoned that mature, quiescent *Drosophila* oocytes, which must exclusively utilize stored mRNAs because transcription is absent, might provide a powerful system for studying FMR1 function in a physiologically-relevant context.

*Drosophila* oogenesis is highly amenable to studying completed, quiescent follicles, since follicles develop sequentially in ovarioles that under appropriate conditions store up to two mature follicles for several weeks. New follicles are continuously produced from stem cells and develop up to stage 8, at which point they are either recycled or allowed to mature if space for another mature follicle is available in the ovariole and dietary nutrients are adequate (Fig. 1A). We found that providing newly eclosed female flies with protein-rich yeast paste for only one day in the absence of mating caused a single wave of two mature follicles per ovariole to develop past the stage 8 checkpoint and go into storage as quiescent follicles (Fig 1B). These finished follicles are maintained in the ovary indefinitely, although a minority slowly turn over and are resorbed or laid. Crucially, no new mature follicles developed from stage 8 follicles for at least two weeks (Fig 1B). To test the developmental competence of the stored oocytes at any age, males can be provided, causing stored oocytes to be ovulated and fertilized, after which they are scored for successful embryogenesis as indicated by hatching into larvae. This protocol allowed us to study mature follicles that had been stored in the quiescent state within the ovary of a living fly for a known period of between one day and two weeks.

**Figure 1.**
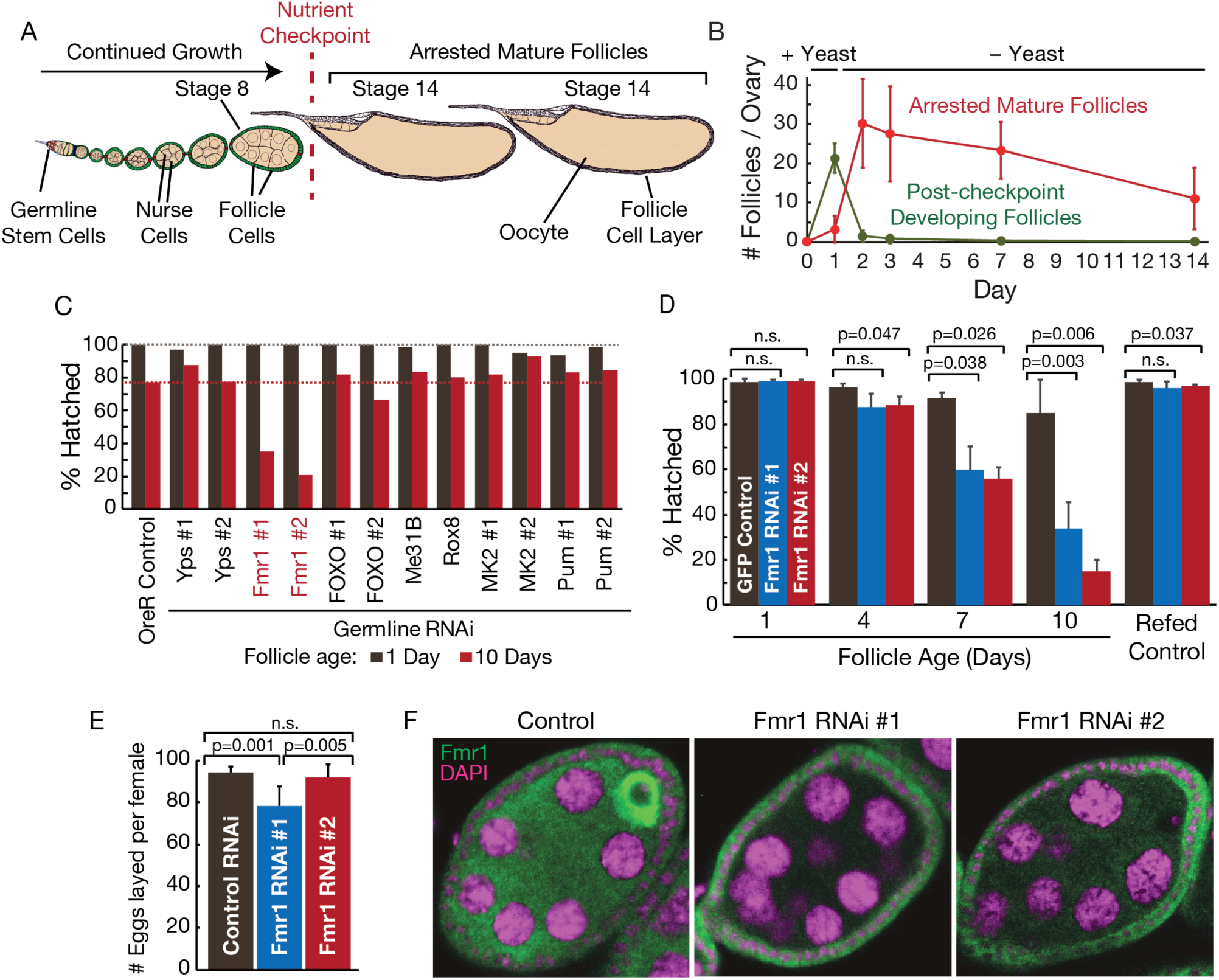
Fmr1 is specifically required during the storage of mature, quiescent stage 14 oocytes in the ovary. (**A**) Schematic of a *Drosophila* ovariole with two stored mature stage 14 follicles. Younger follicles continually develop from stem cells but turn over at the stage 8 nutrient checkpoint unless ovulation of a stage 14 follicle makes space available. Each ovary contains 15 ovarioles. (**B**) Plot shows the stability in the ovary of arrested mature follicles (red) and absence of new maturing follicles (green) following one day of feeding with yeast paste as described here. (**C**) Fmr1 knockdown (line 1 and line 2), but not controls or other gene knockdowns (identified below), specifically reduces 10-day stored (red) but not fresh (black) oocytes from developing into hatching larvae. (**D**) Fmr1 germline RNAi during storage progressively reduces the fraction of mature oocytes competent after 1, 4, 7 or 10 days of storage to support development. Refeeding females to promote maturation of fresh stage 8 follicles restores full developmental potential. (**E**) Fmr1 germline RNAi line 2 does not reduce egg production in the absence of storage. A small reduction in line 1 was due to an off-target effect (see Fig S1D). (**F**) FMR1 antibody staining of stage 8 follicles shows strong FMR1 accumulation in control oocytes, but effective germline-specific knockdown by both RNAi lines.

We tested the function of specific genes during oocyte storage by depleting their transcripts using germline-specific GAL4-driven RNAi, which is produced throughout oogenesis starting in the germline stem cell. Genes required specifically for follicle storage could be recognized because their knockdown follicles would lose developmental capacity more rapidly than wild type follicles during a period of prolonged storage, but would not differ from controls without extended storage. Consequently, we depleted Fmr1 mRNA or several other candidate gene transcripts in oocytes during a storage period of one day or ten days (Fig. 1C). In most gene knockdown lines and in wild type controls the hatch rate was nearly 100% after one day of storage, and only dropped slightly to 80% after 10 days of storage (Fig 1C). In contrast, Fmr1-depleted oocytes generated embryos that hatched normally after one day of storage, but developed only 20-30% of the time following ten days of storage (Fig 1C). Germline Fmr1 RNAi drastically reduced Fmr1 mRNA levels (Fig. S1), and antibody staining confirmed that FMR1 protein was effectively depleted throughout oogenesis specifically in germ cells but not in somatic cells (Fig. 1F).

We validated the Fmr1 requirement of quiescent, stored oocytes by asking if oocyte viability drops continuously over time without Fmr1 (Fig. 1D). The hatch rates of eggs derived from Fmr1 RNAi follicles on day 1 were comparable to controls at 100%, and fecundity was little affected without storage (Fig. 1E). However, the hatch rate fell progressively after storage for 4, 7 and 10 days (Fig 1D). Importantly, the hatch rate after ten days could be fully restored by re-feeding the mother for three days, which caused the remaining stored follicles to be laid, and new mature follicles which had never been stored to develop from stage 8 for testing (Fig 1D). Thus, germline Fmr1 RNAi under the conditions used does not significantly impair stem cells or general follicle development, or compromise hormonal or other non-autonomous aspects of female reproductive function. This differs from Fmr1 mutants which lose stem cells and produce fewer follicles (*19*, *24*), presumably because germline RNAi does not block Fmr1 function in niche cells and other somatic cells that cause these effects. Thus, germline Fmr1 RNAi generates finished oocytes lacking FMR1 as they begin storage, which shortens oocyte viability in the ovary.

We analyzed embryos derived from control and FMR1-deficient oocytes to investigate why FMR1 depletion causes oocytes to prematurely lose their developmental potential. Embryos from control oocytes developed normal nervous systems regardless of prior storage, as shown by staining using a broadly expressed neural marker (Fig. 2A, B). The same was true of embryos derived from Fmr1 RNAi oocytes after one day of storage (Fig. 2C). In contrast, more than 50% of embryos derived from FMR1-deficient oocytes stored in the ovary for ten days developed a severely abnormal nervous system (Fig. 2D, H). While 100% of embryos derived from aged control oocytes appeared normal (Fig. 2E), ventral nerve cord-specific labeling showed missing commissures and breaks in the longitudinal connectives in embryos derived from follicles lacking FMR1 during ten days of storage (Fig. 2F). Embryos with perturbed nervous system development appeared developmentally normal in other aspects, for example undergoing dorsal closure in a manner similar to wild-type embryos (Fig. S2), indicating that the neuronal defects were not due to a generalized deterioration in embryonic function. The severe neural defects were also not just due to an accelerated rate of generalized oocyte deterioration caused by Fmr1 knockdown. We never observed comparable neural defects in embryos derived from normal eggs even when they were stored for longer periods until only 22% of embryos developed to hatching (Fig 2G). Somehow reducing oocyte FMR1 function during egg storage specifically and massively increases the probability an embryo derived from the oocyte will develop with a compromised nervous system.

**Figure 2.**
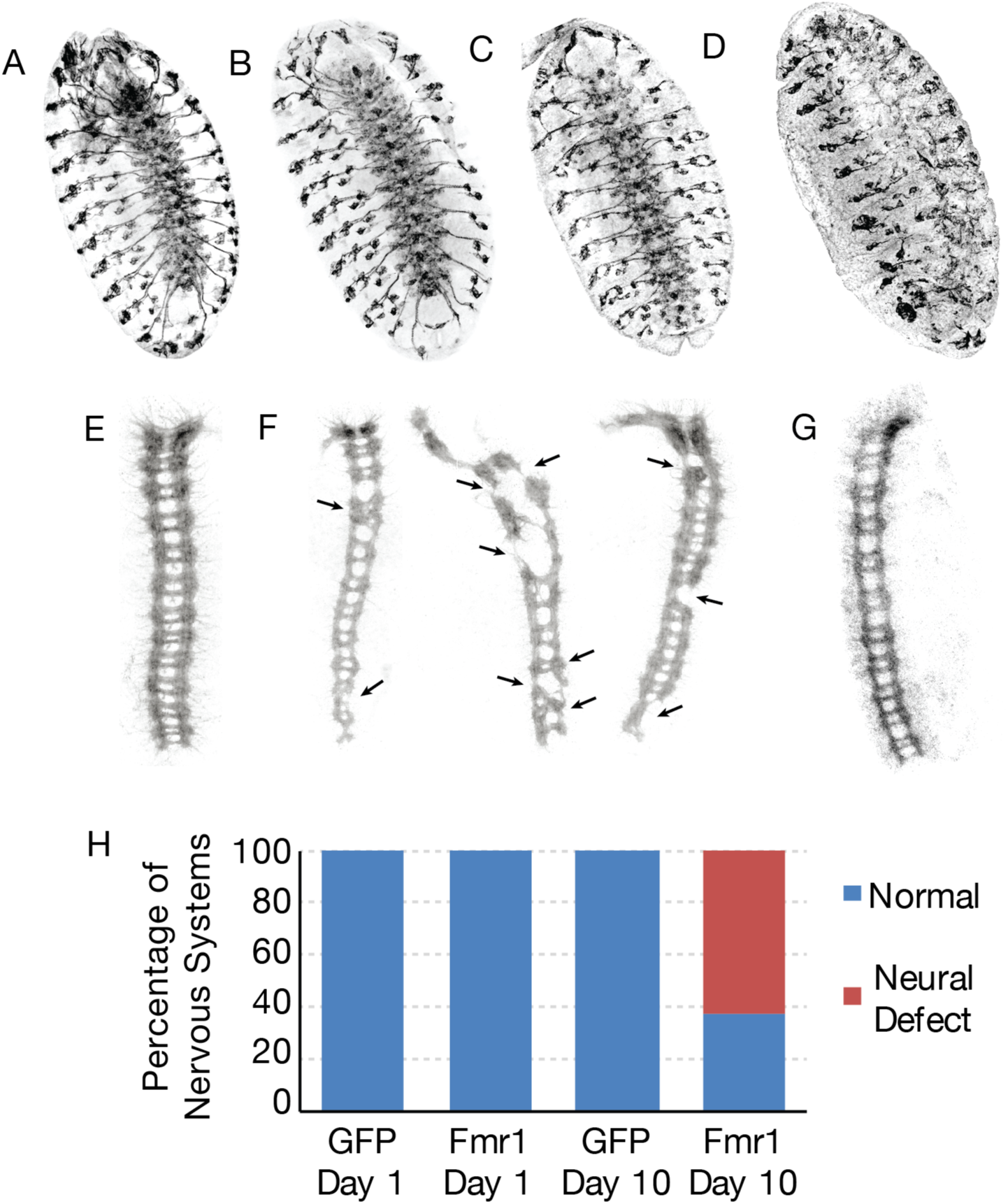
Stored FMR1-depleted oocytes frequently generate embryos with neural defects. (A-D) Staining neurons with 22C10 antibody reveals the normal nervous system of an embryo from a wild oocyte stored 1 day **(A**), wildtype oocyte stored 10 days **(B**), or Fmr1 RNAi oocyte stored 1 day **(C**), and the fragmented, abnormal nervous system from an Fmr1 RNAi oocyte stored 10 days **(D**). **(E-G)** Ventral nerve cords (BP102 staining) show normal structure in embryo developed from wild type oocyte **(E**), but abnormalities including broken connectives (arrowheads) in embryos from stored oocytes lacking FMR1 **(F**) Normal nervous system from a wild type oocyte aged 14 days, when only 22% of derived embryos hatched. **(G)** Percentage of normal or defective nervous systems in embryos from oocytes expressing RNAi to GFP (control) or Fmr1 and held in the ovary until the indicated day.

In order to investigate how Fmr1 maintains stored follicles and preserves their ability to generate a normal nervous system, we examined translation in wild type and FMR1-depleted ovarian tissue. We developed a ribosome profiling protocol based on (*25*) as described in supplemental methods to directly quantify Fmr1-dependent changes in translation in an unbiased, genome-wide manner. Flies were treated to induce the storage of mature follicles and analyzed after 1-2 days in order to identify initial translational changes in healthy Fmr1 knockdown oocytes prior to a reduction in their viability. Although it was necessary to analyze whole ovaries to get enough material for profiling, most of the analyzed ribosomes should still derive from the two stored stage 14 follicles, compared to the much smaller stage 8 and earlier follicles in each ovariole. Interestingly, the ribosome footprints and transcript levels of the vast majority of mRNAs were unaffected by the knockdown of germline-Fmr1 (Fig 3A, B), inconsistent with the idea FMR1 plays a general role in translational control or mRNA stability during storage.

**Figure 3.**
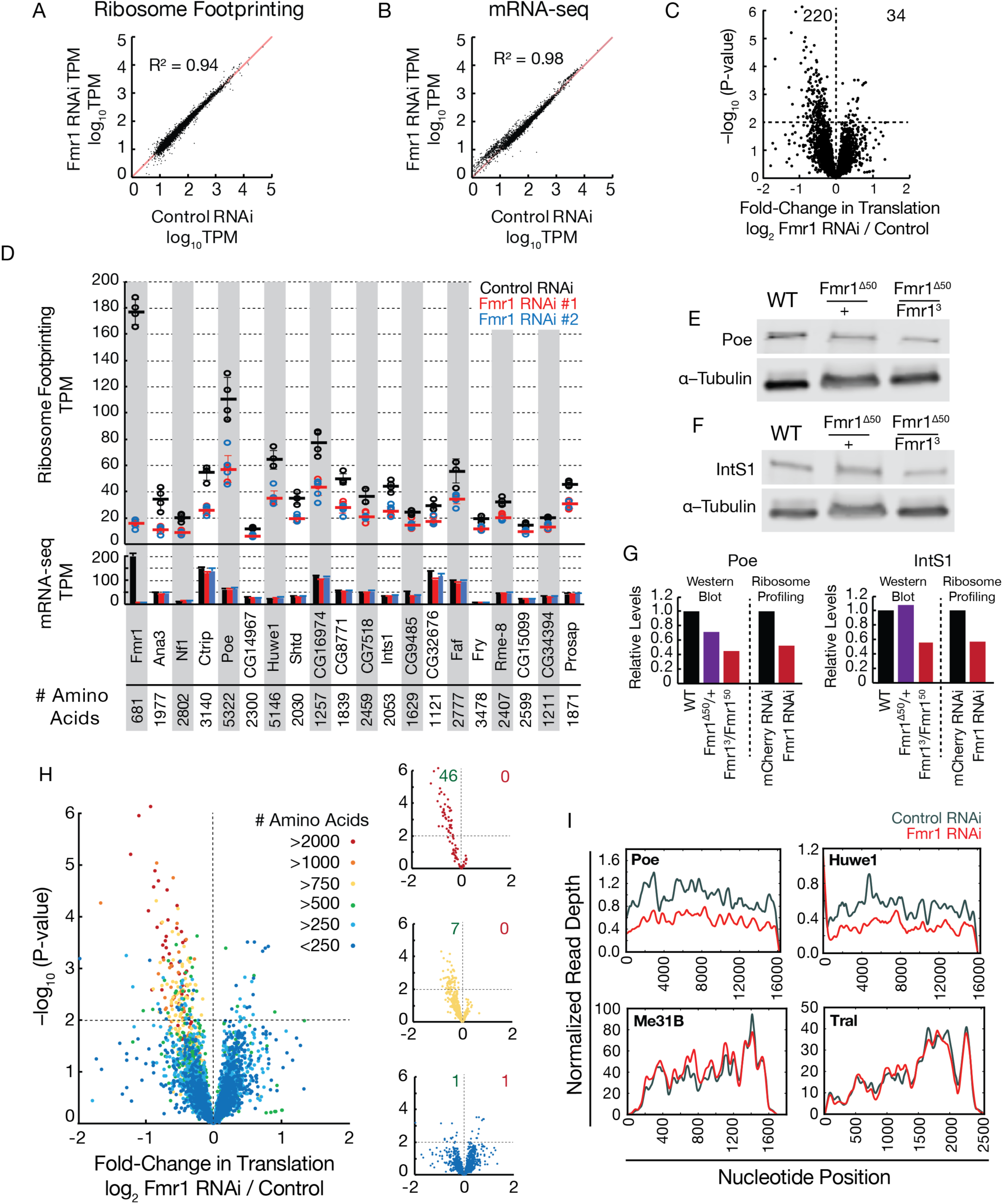
FMR1 stimulates the translation during storage of transcripts from multiple intellectual disability and autism genes. **(A)** Absence of significant global change in ribosome footprinting levels (log_10_TPM; transcripts per million) between control vs Fmr1 RNAi oocytes after 1-2 days of storage. **(B)** Absence of significant global change in mRNA levels determined by RNA-seq (log_10_TPM) between control vs Fmr1 RNAi oocytes after 1-2 days of storage. **(C)** Identification of 220 significant translational targets repressed by Fmr1 RNAi and 34 significant translational targets upregulated by Fmr1 RNAi (p value < 0.01) as measured by ribosome footprinting. **(D)** Ribosome footprinting TPM (above) and RNA-seq TPM (below) for the 40 downregulated genes with the lowest p-values (sorted by fraction reduced FMR1/control). Gene name and protein size is shown below and the values for controls (N=4), RNAi #1 (N=3) RNAi #2 (N=4) are plotted, with means and SDs shown. **(E-G)** Quantitative Western blots for Poe (**E**) and IntS1 (**F**) using isolated mature oocytes from the indicated Fmr1 genotypes. **(G)** Comparison of relative protein levels for indicated genotypes determined by Western blotting and ribosome footprinting shows close agreement. **(H)** FMR1 preferentially augments the translation of large proteins. Proteins >2000, 750-1000, or <250 amino acids are shown at right for clarity. Numbers in green and red indicate percentage of genes downregulated or upregulated respectively (p < .01). **(I)** FMR1 does not impair translational processivity. Normalized read depth is plotted for two FMR1 targets (Poe and Huwe1) and two non-targets (Me31B and Tral). In Fmr1 RNAi oocytes, footprints are reduced at all positions along the mRNA.

We identified statistically significant oocyte target genes by analyzing seven independent ribosome profiling measurements of FMR1 germline RNAi (using both line 1 and line 2) and four independent measurements of control profiles (Fig. 3C). 220 oocyte-expressed genes, including Fmr1, were significantly reduced in expression and 34 genes showed more expression in FMR1 RNAi compared to control (Table S1, S2). Except for Fmr1 itself, the significantly altered genes changed relatively little, only 1.5-2.5-fold. However, an exceptionally high fraction of the downregulated genes, at least 37/219, are orthologs of human genes that have been implicated in human neuro-developmental syndromes (Table S3).

The twenty most significantly downregulated oocyte genes include many well known genes associated with human intellectual disability syndromes and autism (Fig. 3D). For example, the neurofibromatosis gene, Nf1, which is associated with cognitive and behavioral disorders and neural tumors (*26*, *27*), was reduced 2.3-fold. Ctrip, encoding a HECT-domain E3 ubiquitin ligase whose human homolog Trip12 is associated with intellectual disability and autism (*28*, *29*), was reduced 2.1-fold. The gene encoding the N-end rule E3 ligase adaptor POE was reduced 1.9-fold; Poe’s human homolog Ubr4 was recently implicated in early onset dementia (*30*). The production of HUWE1, another HECT-domain E3 ubiquitin ligase linked to idiopathic intellectual disability and schizophrenia (*31*, *32*) was reduced 1.9fold. Prosap, encoding a postsynaptic density scaffolding protein, was reduced 1.5-fold. Its human orthologs Shank 1/2/3, are associated with autism, intellectual disability, and schizophrenia (*33*, *34*). Moreover, homologs of several other dominant autism/disability genes were also substantially reduced, including Arid1B, Rere, Brwd3, Med12, Btaf1, Depdc5, Hdc, YY1, and Cyfip1 (Table S2). Since mutations in these genes are dominant (*35*), a two-fold reduction in expression is potentially significant, even for a single target.

Several other genes in Fig. 3D, Ana3/Rttn, IntS1, Shtd/Anapc1, and Faf/Usp9Y have been linked to human recessive neurodevelopmental syndromes and autism (*35*). Homologs of many additional human genes associated with recessive neural defects were also significantly reduced, including Anapc1, Dmxl2, Brd4, Mpp6, Mkl2, Stxbp5, Pi3k3g, Atrx, Phrf1, Ep400, Sbf1, Usp45, Taf1, and Tbc1d31 (Table S3). The reduced production during oocyte storage in the absence of full FMR1 function of so many genes important for nervous system development provides a highly plausible explanation for why stored FMR1 RNAi oocytes lose the ability to support embryonic nervous system development.

To verify that our ribosome profiling measurements identify genes whose translation is actually reduced in Fmr1 RNAi oocytes, we generated polyclonal anti-POE antibodies and obtained an antibody specific for INTS1. We carried out quantitative Western blotting (Fig. 3E, F), and calculated that both proteins showed reductions in stored oocytes lacking FMR1 function closely similar to those determined from ribosome profiling experiments (Figure 3G; Figure S5).

Fmr1 might target this large collection of neurodevelopmental genes specifically, as a sort of master neural regulator. However, our results uncovered a general characteristic shared by most of the proteins altered by Fmr1 RNAi that provides a simpler explanation. Almost all of the affected proteins are much larger than the average *Drosophila* protein (Fig 3D). Whereas only 2.7% of all *Drosophila* proteins contain 1750 or more amino acids, more than half (21/37) of the genes in Table S3 are at least this size (p-value 4.4 x 10^−89^, Chi-squared test).

Moreover, there was a striking correlation between protein size generally and a requirement for FMR1 during storage (Fig 3H, Fig. S3A, B). At least 40% of proteins with greater than 2,000 amino acids showed a significant FMR1 requirement, and the entire fold change plot was strongly skewed even for smaller proteins 750-1000 amino acids (Fig 3H, Fig. S3A, B). In contrast, protein abundance was uncorrelated; FMR1 stimulates mRNAs across all expression levels (Fig. S3C, D). The human homologs of the neurodevelopmental genes that require FMR1 are also exceptionally large, and strongly correlated in size to their *Drosophila* counterparts (Fig. S6). Conversely, the few genes upregulated in FMR1 RNAi were all quite small (Fig. S4A, B), and their homologs were not associated with neurodevelopmental disease genes.

Large stored mRNAs might be harder to translate if processivity is reduced. To test for reduced processivity, we examined the distribution of ribosome footprints across the length of several large mRNAs whose translation is reduced in FMR1 RNAi. Contrary to the prediction of the processivity model that footprints would show little difference at the amino terminus but decline sharply toward the C-terminus, we observed a uniform reduction in footprints across the entire coding sequence (Fig 3I). We found that the length of a transcript’s coding sequence contributed most strongly to a translational requirement for Fmr1, along with smaller contributions from the 5’ and 3’ untranslated regions (Fig S4C, D).

Since FMR1 affects the translation of 37 genes important for neural development based largely on a general characteristic (size), and since each target is reduced by a relatively small amount, one might expect that the effect of any given target gene would be incremental. To test if an individual target gene might itself be functionally required to maintain *Drosophila* oocytes in storage, we selected the Poe/Ubr4 gene encoding the largest putative Fmr1 target. Poe mutant *Drosophila* share multiple phenotypes with Fmr1 mutants, including male sterility, inability to fly, and increased neuromuscular junction synaptic excitability (*8*, *36*, *37*). We compared the hatching frequency of eggs from Poe or Fmr1-deficient females following various periods of follicular storage, and observed that stored Poe mutant oocytes lost developmental competence at the same rate or even slightly faster than Fmr1 germline RNAi oocytes (Fig 4A). The hatch rate of embryos derived from unstored (1 day) Poe mutant follicles was comparable to controls, demonstrating that Poe, like Fmr1, plays a functionally important and specific role in maintaining ovarian follicles during developmental arrest.

**Figure 4.**
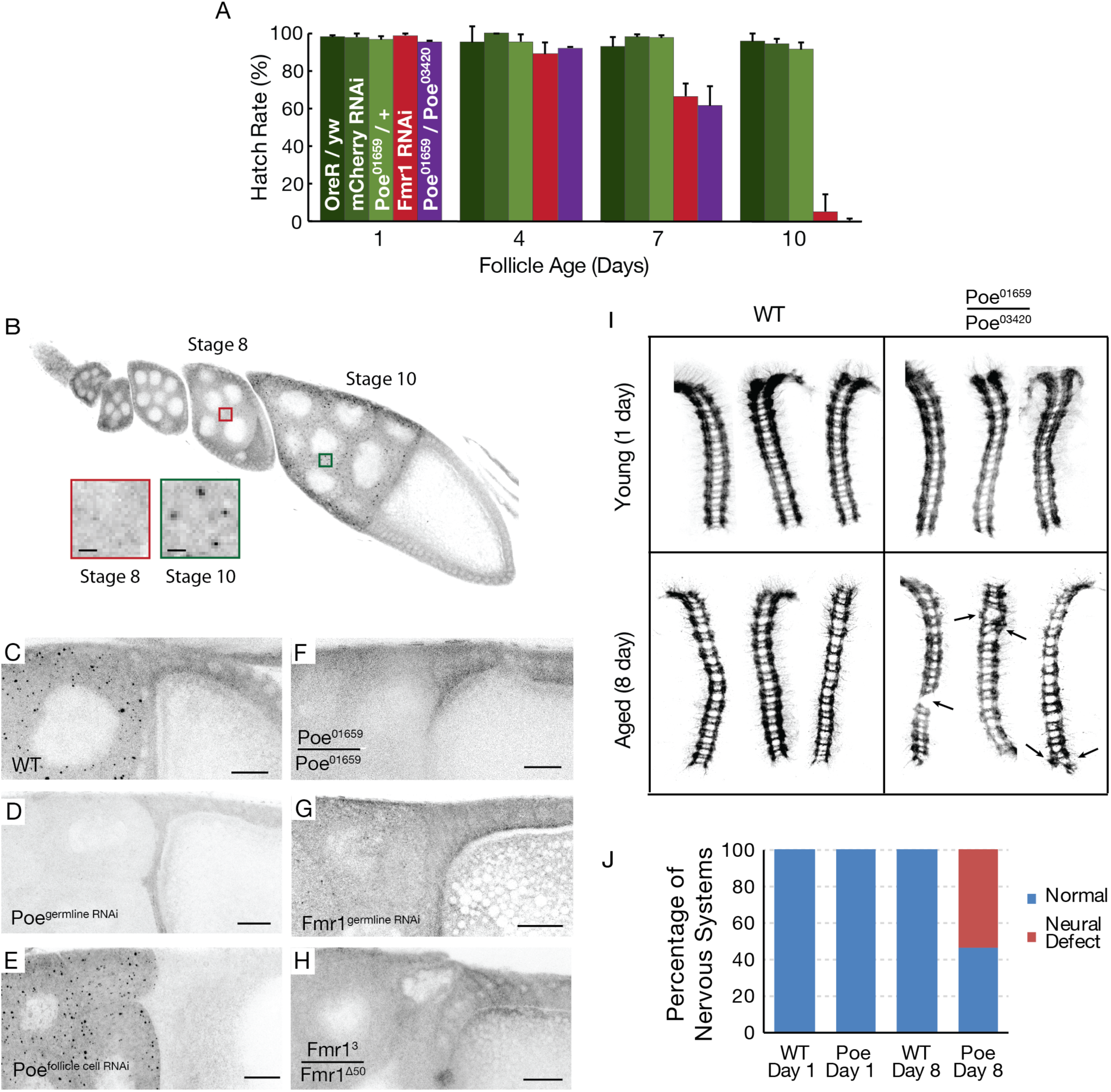
The Fmr1 target Poe/Ubr4 is itself required for oocyte storage and neural development. (**A**) Poe mutation accelerates oocyte decline during storage as much as Fmr1 RNAi. Hatch rate as a function of days in storage (follicle age) prior to fertilization and testing for development. (**B**) POE forms germline granules in maturing follicles. An ovariole stained with anti-Poe antibodies reveals generalized cytosolic germline staining up until stage 8. By stage 10, Poe is expressed in cytoplasmic granules (inset). Bar = 3μm. (**C-H**) Part of a stage 10 follicle is shown after staining with anti-Poe antibodies. Many Poe granules are seen in wild type (**C**), but not in Poe^germline^ ^RNAi^ (**D**) or Poe^01659^ homozygotes (**F**), whereas Poe^follicle^ ^cell^ ^RNAi^ is like the control (**E**). Germline-specific reduction of Fmr1 (Fmr1^germline^ ^RNAi^) (**G**), or Fmr1-null mutation (**H**) eliminates Poe granule staining, validating Poe as an Fmr1 target in vivo. (**I-J**) Poe is required for stored oocytes to support neural development. Nerve cords from embryos developed from oocytes of the indicated genotypes and storage times are shown. Like Fmr1, oocytes require Poe during storage to maintain the ability to form an intact nervous system. Arrowheads show gaps in the nerve cords of embryos from stored Poe^01659^ homozygous oocytes. (**J**) Chart showing the percentage of embryos with intact or defective nervous systems that developed from wild type (WT) or Poe mutant (Poe) mature oocytes stored for the indicated number of days.

We analyzed the expression pattern of POE during follicle development, using our polyclonal antibody. POE protein was largely cytosolic during early oogenesis, and was strongly induced after the stage 8 checkpoint as follicles approach their final size. Beginning in nearly mature follicles, POE became associated with 0.5-2 micron spherical particles in germ cells (Fig. 4B). The observed staining, including the specific particles, was not detected in ovaries expressing Poe germline RNAi but was unaffected by follicle cell-specific RNAi (Fig 4C-E). POE particles were also absent in a Poe mutant which has a large reduction in Poe expression (Figure 4F). As predicted (Fig. 3D), POE protein levels were reduced in Fmr1-germline RNAi and Fmr1-null egg chambers, and POE particles were drastically reduced or eliminated from Fmr1-germline RNAi and Fmr1-null follicles, respectively (Figure 4G, H). These data indicate that Poe is essential to maintain mature ovarian follicles in an arrested state, and that Fmr1 functions to induce POE accumulation and particle formation starting just prior to follicle completion.

When we analyzed the ventral nerve cords of embryos developing from Poe mutant females whose oocytes had been stored for 1 day, they were normal (Fig. 4I, J). Remarkably, more than 50% of embryos developing from 8 day-aged oocytes from these same animals had a disrupted nervous system, and showed similar defects in neural commissures and connectives as in aged Fmr1 RNAi oocytes (Fig. 4I, J). These observations demonstrate that Poe is a major functional target of Fmr1, and that Poe is itself essential to maintain oocyte viability and neurodevelopmental competence during storage.

Previous attempts to identify Fmr1 targets largely relied on proximity-based strategies to identify cross-linked mRNAs followed by immunoprecipitation of FMR1 (*21*-*23*). It is not clear that the cells used in these studies were highly enriched in FMR1-containing granules engaged in translational control. Moreover, RNPs heavily concentrate mRNAs and RNA-binding proteins into dynamic liquid-like particles, potentially increasing the probability of spurious, non-functional interactions in cross-linking studies. Despite these potential difficulties, the 842 putative Fmr1 targets identified (*21*) were significantly enriched in our study. These targets comprise 6.9% of ovary-expressed *Drosophila* genes (Table S4) with human homologs, but 21.5% of the 219 Fmr1 target genes in Tables S1 (p<3.3E-16, Chi squared test). Moreover, 51% (19 /37; p < 1.1E-24, Chi squared test) of the autism-associated genes identified by these authors were among the autism-related genes significantly affected in our assays (Table S3). Thus, previous studies identified a subset of the targets seen using our approach.

A major difference with most previous studies is that FMR1 has generally been regarded as a translational repressor (*21*, *38*, *39*), rather than a translational activator, as seen here. There have been some previous reports of FMR1 acting as a translational activator (*23*, *40*, *41*). Indeed, the common association of FMR1 with translational repression may be indirect. E3 ubiquitin ligases are often large proteins, and at least six are positively regulated by Fmr1 (Table S1). Poe/Ubr4 in particular is a member of the N-end rule family of N-recognins, which catalyzes the degradation of a broad range of substrates (*42*). The loss of Poe/Ubr4 activity in FMR1-deficient cells would likely cause Poe/Ubr4 targets to be over-expressed. The phenotypic similarities between Poe/Ubr4 and Fmr1 mutants, including specific synaptic defects (*8*, *36*), and in egg storage, suggest that Poe/Ubr4 may be a functionally important Fmr1 target.

Why do the effects of FMR1 RNAi or Poe mutation on egg viability and neural development only occur after storage? Our experiments show that FMR1 RNAi already abolishes FMR1 protein (Fig. 1F) and Poe expression (Fig. 4G) before oocytes enter storage. The target genes we identified are already reduced within 1-2 days of storage, before oocytes lose the ability to develop normally. We propose that FMR1-deficient follicles fail because the levels of some or all of FMR1 target proteins decline further below critical levels over time during storage, and/or because secondary effects caused by the decreased expression of targets are compounded over time. For example, a decrease in levels of Poe/Ubr4 and other target E3 ligases might lead to over-accumulation of normal or aggregated proteins that compromise neural development.

The experiments reported here provide new insight into the functional role of Fmr1 in the ovary and during neural development as a size-dependent positive translator regulator. Mammalian and yeast mRNAs associate on the basis of size with stress granules where their translation is repressed (*43*). FMR1 may function to counteract this inherent tendency of large mRNAs to be segregated into inactive RNP particles. Alternatively, Fmr1 might directly or indirectly promote translation initiation in association with RNPs, or it might affect mRNA transport along microtubules to sites of active initiation (*44*, *45*).

Our work also suggests why FMR1 mutations alter mammalian ovarian follicle development and maintenance. Reductions in FMR1 levels are intrinsically deleterious to stored mature *Drosophila* follicles and shorten their functional lifetime. The larger ovarian size and elevated antral follicle content of FMR1-null mice (*22*) may reflect similar defects in stored, mature follicles that block their selection for ovulation and cause them to accumulate. Specific human alleles associated with ovarian decline (*46*, *47*) may exert a dominant negative effect that compromises stored primordial follicles as well as stored mature follicles, leading to primary ovarian insufficiency. At present, evidence is lacking that defects which may arise during storage of mature mammalian oocytes contributes later to neural defects like in Drosophila, but this would be worthy of investigation.

In conclusion, the discovery that FMR1 is needed to maintain the *Drosophila* oocyte’s competence for neural development by boosting the translation of multiple large proteins that contribute to neural and cellular function provides fundamental new insight into development and senescence. We propose that this represents a general function for FMR1 in cells that rely heavily on translational control mechanisms, such as oocytes and neurons. Improved knowledge of how FMR1 works and the identities of potentially important targets open new opportunities to monitor susceptible cells, and to intervene to mitigate declining levels of the most critical targets. Small molecule agents that counteract the tendency of large mRNAs to be segregated into inactive granules represent a potentially valuable therapeutic approach in Fragile X patients. The possibility that key large proteins become reduced in other neurodevelopmental syndromes should also be considered. In particular, our data suggest that Poe/Ubr4 ranks as an important target whose role in disease should be further investigated. Finally, the likelihood that adult neurons use the pathways of translational regulation assayed here suggests that under-production of critical large proteins contributes to some adult-onset neural impairments such as schizophrenia and dementia. Continued study of these highly conserved pathways in *Drosophila* represents a powerful and efficient means to address many of these issues.

## Acknowledgements

We thank Allison Pinder for help generating RNA-seq libraries. We are grateful to Joshua Dunn (UCSF) for technical advice which greatly enabled our ribosome footprinting experiments. We thank Eric Wagner (UTMB) and Thomas Jongens (UPenn) for generously providing the IntS1 antibody and the Fmr1^3^ mutant fly strain respectively. This work was supported by funding through the Jane Coffin Childs Memorial Fund (E.G.) and through the generous support of the Howard Hughes Medical Institute (A.C.S.).

## Supplementary Materials

### Fly Stocks

The following *Drosophila* stocks obtained from Bloomington *Drosophila* Stock Center were used in this study: UAS-Fmr1 RNAi #1 (Harvard TRIP Collection, HMS00248), UAS-Fmr1 RNAi #2 (Harvard TRIP Collection, GL00075), UAS-mCherry RNAi (Harvard TRIP Collection, VALIUM20-mCherry), UAS-GFP (Harvard TRIP Collection, UAS-GFP.VALIUM10), Germline Gal4 (nos-Gal4VP16, third chromosome insertion), Follicle Cell Gal4 (Janelia Gal4 Collection, 10H05), and Fmr1▵^50^. We obtained the Fmr1^3^ stock from the laboratory of Thomas Jongens (UPenn).

### Follicle Survival Assay

Virgin female flies were collected over 8 hours at room temperature or 16 hours at 18C and were supplied with wet yeast paste (dry yeast + water to the consistency of peanut butter) for 24 hours. Fattened flies were then transferred to molasses vials or plates for the remainder of the experiment (180mL molasses, 44g agar, 37 mL 5% tegosept, 1112 mL water). Egg laying was induced by adding 10 3-8 day old males to 10 females on scored molasses plates for 24 hours. Prior to addition, males were kept separately from females for 2 days. Egg hatch rates were determined by visual examination.

### *Drosophila* Ovary Immunostaining

Ovaries were dissected in Grace’s buffer and fixed in 4% formaldehyde (37% formaldehyde diluted in PBST (0.2% BSA, 0.5% Triton X-100 in 1XPBS)) for 12 minutes. Ovaries were washed 3 times in PBST for at least 20 mins per wash and incubated overnight with primary antibodies at 4C with gentle agitation on a nutator. Ovaries were then washed 3 times in PBST for at least 20 min per wash and incubated with secondary antibodies for either 2 hours at room temperature or overnight at 4C. Ovaries were then washed 3 times for 2 hours per wash in PBST, adding DAPI (5mg/mL stock diluted 1:20,000 in PBST) into the final wash. Primary antibodies used were anti-dFmr1 6A15 (1:1,000, Abcam) and Poe (1:100). The Poe polyclonal rabbit antibody was generated against a recombinant N-terminal fragment containing residues 40-208 (Proteintech).

### Quantitative Western Blots

Female flies were maintained on wet yeast paste with males for 5 days, and unstored mature follicles were isolated manually by filtration following collagenase treatment of dissected ovaries with 5 mg/mL collagenase (Sigma-Aldrich) in Grace’s Buffer for 10 minutes. Follicles were lysed in sample buffer, and were loaded onto a 4-15% Tris-glycine gradient gel, and electrophoresed at 120V for 1hr. Gels were then transferred in Towbin’s buffer at 350 mA for 4 hours onto a PVDF membrane. Membranes were blocked in Odyssey Blocking Buffer for 30 mins at room temperature, and incubated with primary antibodies against Poe (generated by Proteintech, 1:100), IntS1 (1:2,000, laboratory of Eric Wagner at UTMB), or alpha-Tubulin (1:2,500 DM1A, Millipore) overnight at 4C with gentle agitation in Odyssey Blocking Buffer containing 0.1% Tween 20. Membranes were washed 3 times with TBST and incubated with NIR secondary antibodies (1:15,000 LI-COR) for 1 hour at room temperature. Membranes were washed for 3 times with TBST and twice with TBS for 5 mins per wash, dried, and imaged using an Odyssey CLx imager.

### Ribosome Footprinting and mRNA-seq

Ribosome footprinting was performed as per Dunn and Weissman (*1*) with several modifications. Virgin female flies were collected and placed on wet yeast paste for 24 hours, and then transferred onto molasses vials for either 24 hours or 48 hours. 40 pairs of ovaries were isolated by manual dissection in Grace’s Buffer and flash frozen in liquid nitrogen. Ovaries were lysed in 200uL of lysis buffer (0.5% Triton X-100, 150mM NaCl, 5mM MgCl2, 50 mM Tris, pH 7.5, 1mM DTT, 20ug/ml emetine, 20 U/mL SUPERaseIn, 50 uM GMP-PNP) by grinding for 15 secs followed by trituration through a 26-gauge needle seven times. 10uL of lysate was removed for the preparation of poly(A)-selected, strand-specific mRNA-seq libraries (TrueSeq Kit, Illumina). Ribosome footprints were generated by incubation of lysate with MNase at a ratio of 1U per ug total RNA for 40 minutes at 25C, then the reaction was quenched by adding EGTA to 6.25mM final concentration. Ribosomes were sedimented through a 34% sucrose cushion for 4 hours at 70,000 rpm in a TLA100.4 rotor, and pellets were resuspended in 10mM Tris, pH 7.0. RNA was then extracted from monosomes through phenol/chloroform extraction, and footprints were size-selected on a 15% TBE-urea gel. Dephosphorylation and ligation of the 3’ adapter was performed by incubation of footprints with shrimp alkaline phosphatase (rSAP, NEB) for 1 hr at 37C, and the reaction was quenched by incubation at 65C for 5 mins. The 3’ adapter ligation was performed in the same tube by adding 2uM of a pre-adenylated adapter (rAppTGGAATTCTCGGGTGCCAAGG) with T4 RNA ligase 2, truncated (NEB). Ligated products were then size-selected on a 10% TBE-Urea polyacrylamide gel. Reverse transcription was performed with Superscript III using a primer: /5phos/GATCGTCGGACTGTAGAACTCTGAACGTGTAGATCTCGGTGGTCGC/isp18/ CACTCA/isp18/CCTTGGCACCCGAGAATTCCA and the reaction was quenched by incubation with 0.1M NaOH for 20 minutes at 98 ^o^C. Following rRNA depletion, cDNA libraries were circularized by two sequential reactions with CircLigase and amplified by 10-12 PCR cycles with primers compatible with Illumina flow cells. Alignment of sequencing data was performed as in Ingolia et al 2012 *(1)* and quantified with Cufflinks (*2*) and Plastid (*3*). Data quantified with Plastid were normalized to the sum of read depths covering mRNA sequences. Data quantified with Cufflinks were pre-filtered against *Drosophila* noncoding RNA sequences and were normalized to transcripts per million reads (TPM) following manual removal of follicle cell-specific genes. Genes whose expression was detected by RNA-seq in stage 14 follicles but not in 0-2 hour embryos *(4)*, such as chorion genes, were considered to be follicle cell specific and were excluded from our studies of oocyte expression.

---

**Supplementary Figure 1.**
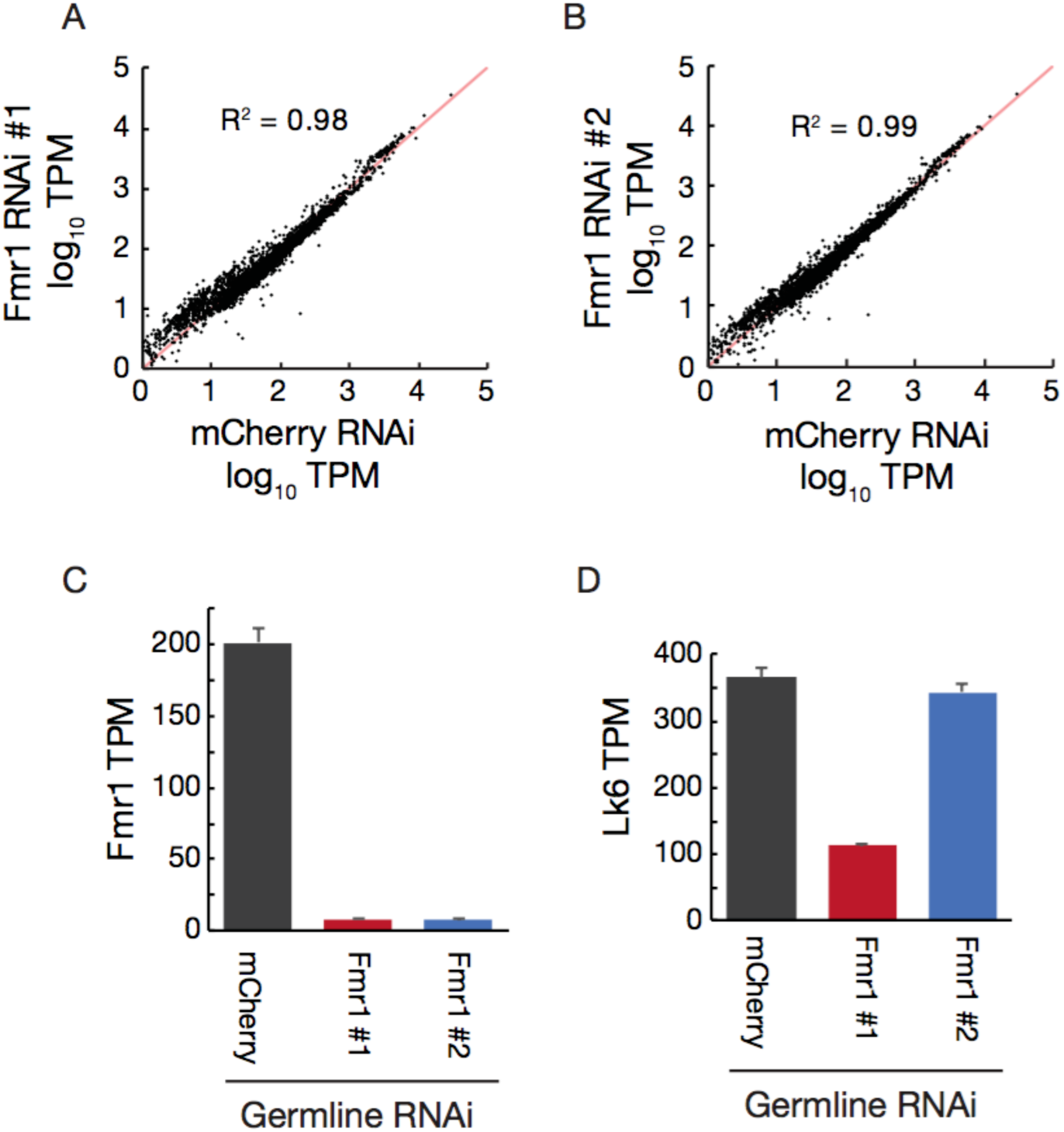
Specificity of Fmr1 RNAi lines. (**A, B**) Absence of global changes of log_10_TPM mRNA levels between control (mCherry) and **(A)** Fmr1 RNAi #1 or **(B)** Fmr1 RNAi #2 as determined by mRNA-seq of oocytes after 2 days of storage. **(C)** Drastic reduction in Fmr1 mRNA levels in Fmr1 RNAi lines as compared to controls as determined by mRNA-seq of oocytes after 2 days of storage. **(D)** Lk6, implicated in growth control *(5)* is a prominent off-target of Fmr1 RNAi #1.

**Supplementary Figure 2.**
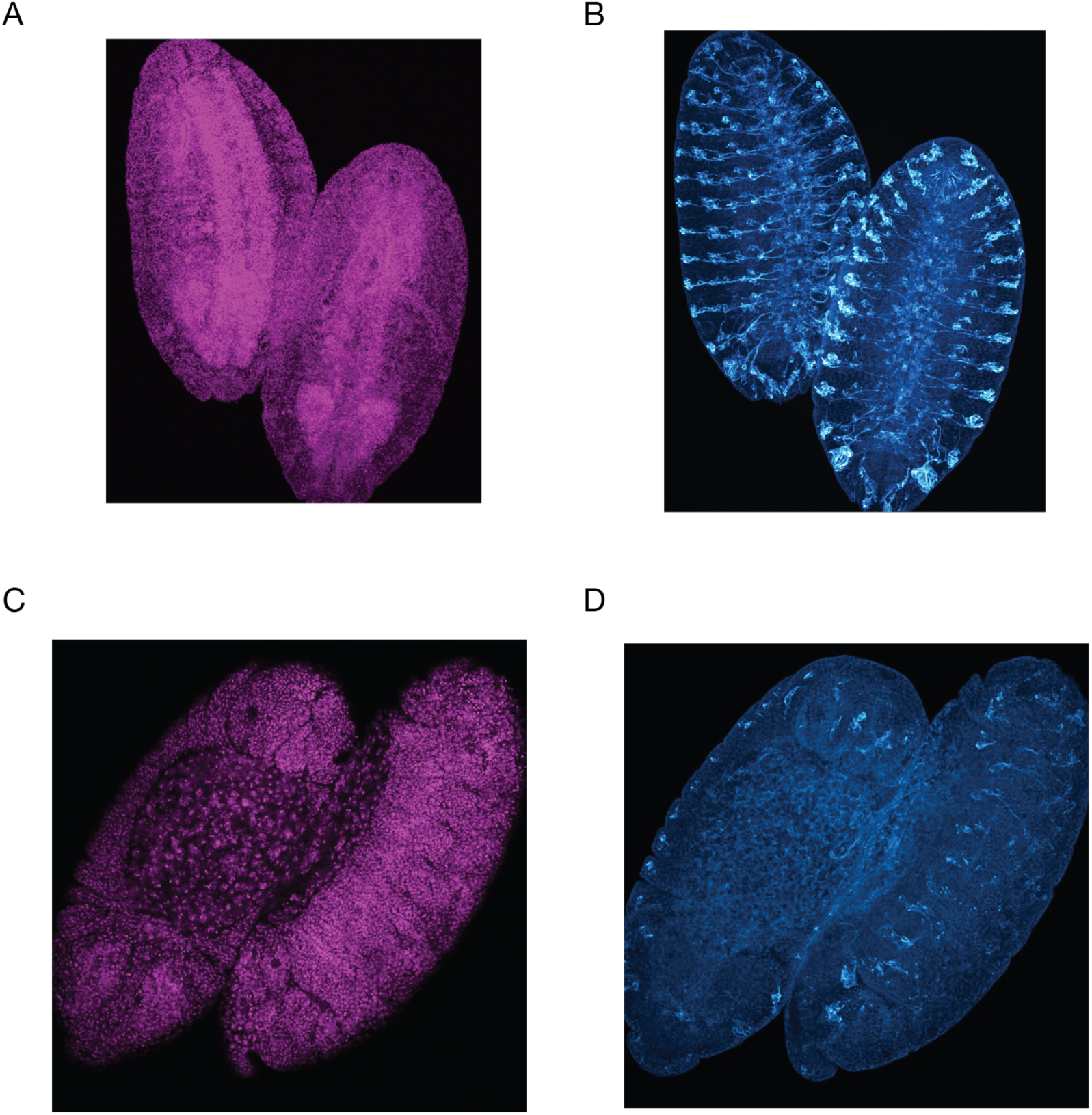
Specificity of neuronal defects in embryos derived from aged FMR1 knockdown oocytes. (**A, B**) Nuclear (DAPI) **(A)** and neuronal (mAb 22C10) **(B)** staining of GFP control 16-20 hour embryos derived from oocytes aged for 10 days. **(C, D)** Embryos derived from equivalently aged FMR1 knockdown oocytes are slightly delayed in their development as compared to controls, but are morphologically normal as judged by nuclear staining **(C),** and exhibit a perturbed nervous system **(D)**.

**Supplementary Figure 3.**
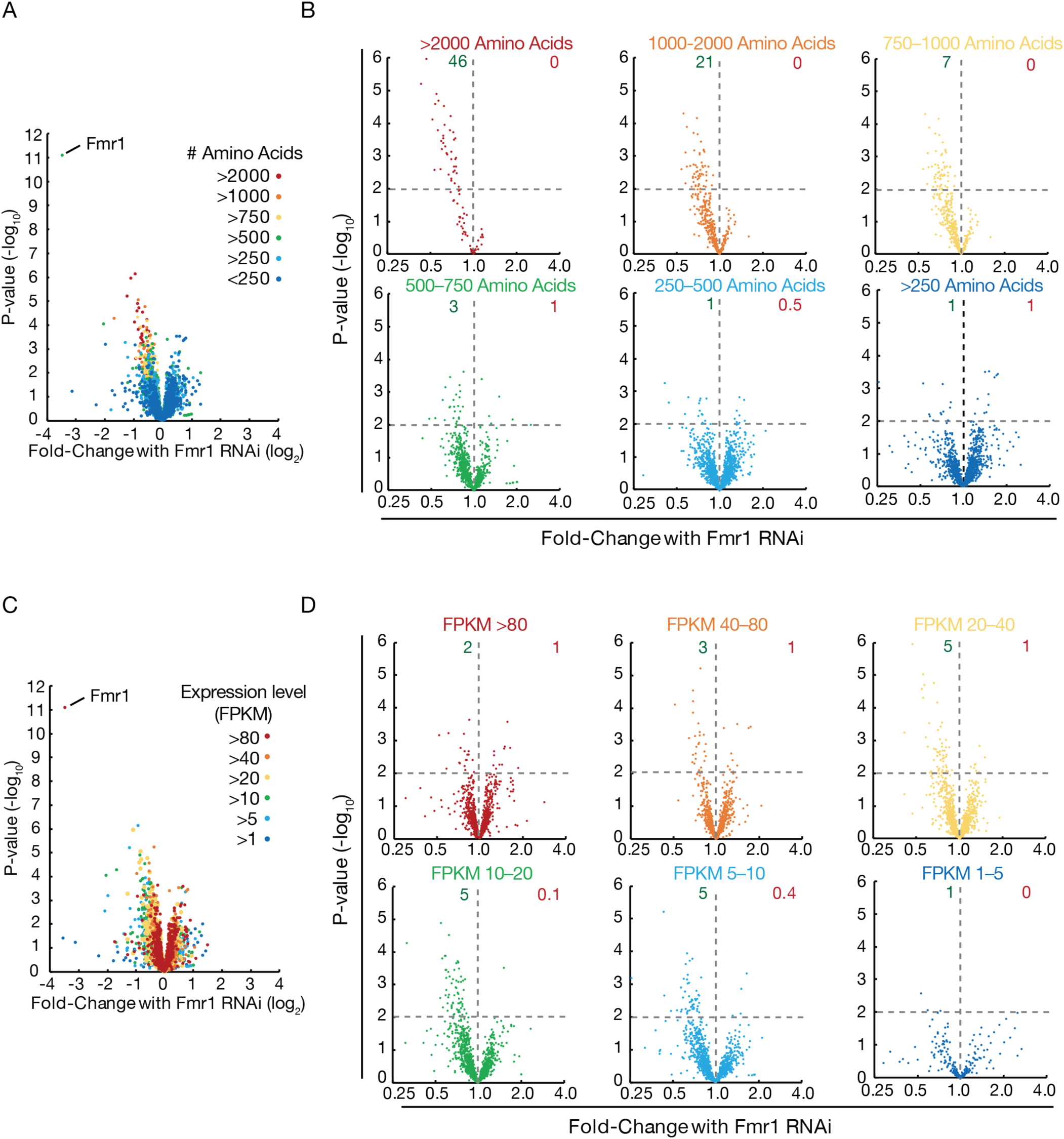
FMR1 is a size-dependent activator of translation. **(A)** FMR1 selectively reduces the translation of transcripts encoding the largest proteins. Zoom-out of volcano plot shown in Fig. 3H to show all transcripts. **(B)** Individual size classes from **(A)** are shown separately for clarity. Numbers in green and red indicate percentage of genes downregulated or upregulated respectively (p < .01). **(C, D)** Translational activation by FMR1 is uncorrelated to expression level. Data are separated by indicated expression level (FPKM value) for clarity.

**Supplementary Figure 4.**
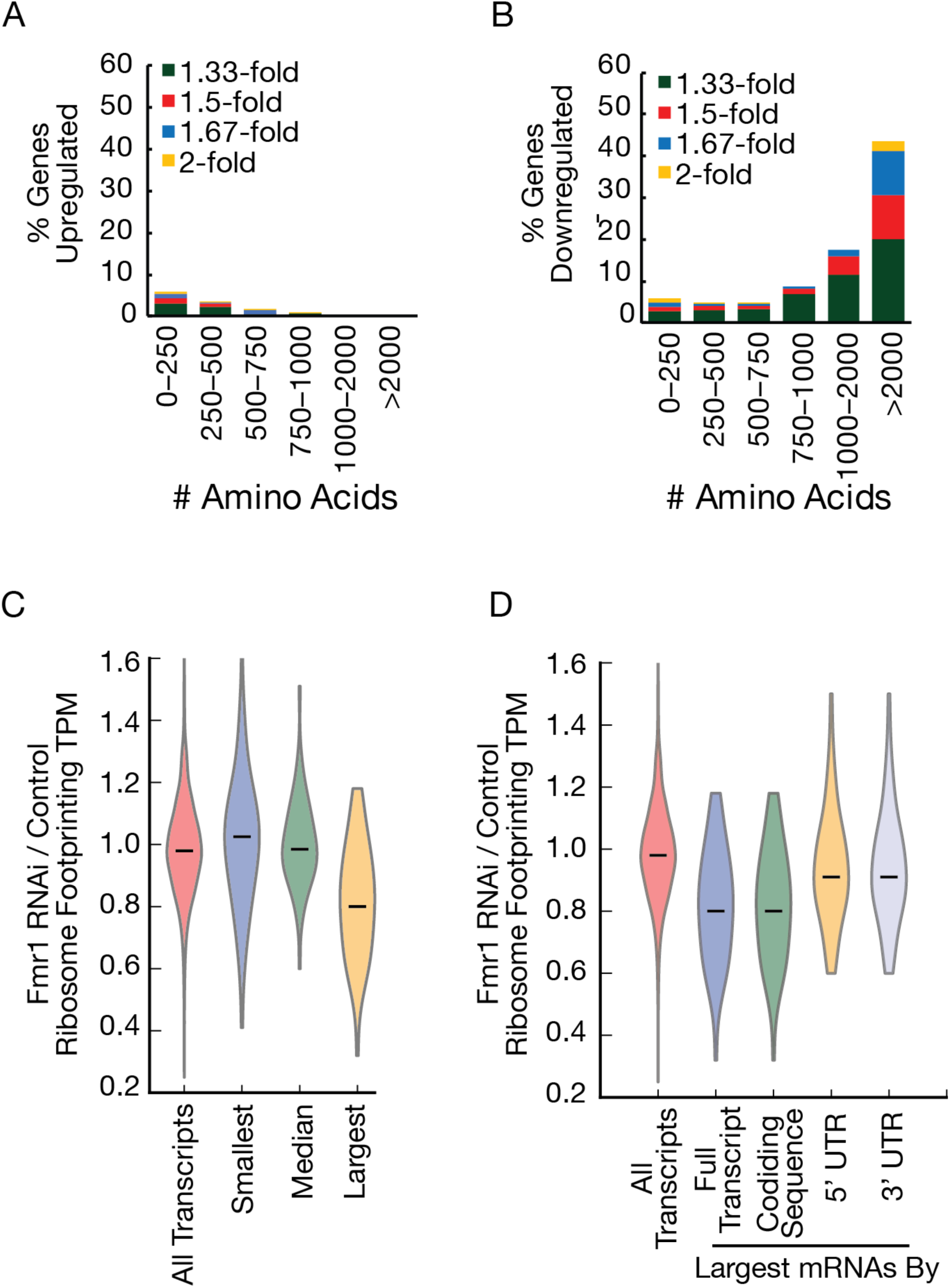
FMR1 selectively enhances the translation of large proteins. **(A, B)** A small fraction of small proteins show increased ribosome footprinting levels in Fmr1 RNAi oocytes as compared to controls after 1-2 days of oocyte storage **(A)**, while a large fraction of large proteins show decreased ribosome footprinting levels (**B). (C)** The longest (top 2%) mRNAs by transcript length show reduced translation in Fmr1 RNAi samples as compared to all transcripts and the smallest/median 2%. **(D)** The longest (top 2%) of mRNAs by transcript and coding sequence length show reduced translation in Fmr1 RNAi samples as compared to smaller effects for mRNAs with the longest 2% of 5’ and 3’ UTR sequences.

**Supplementary Figure 5.**
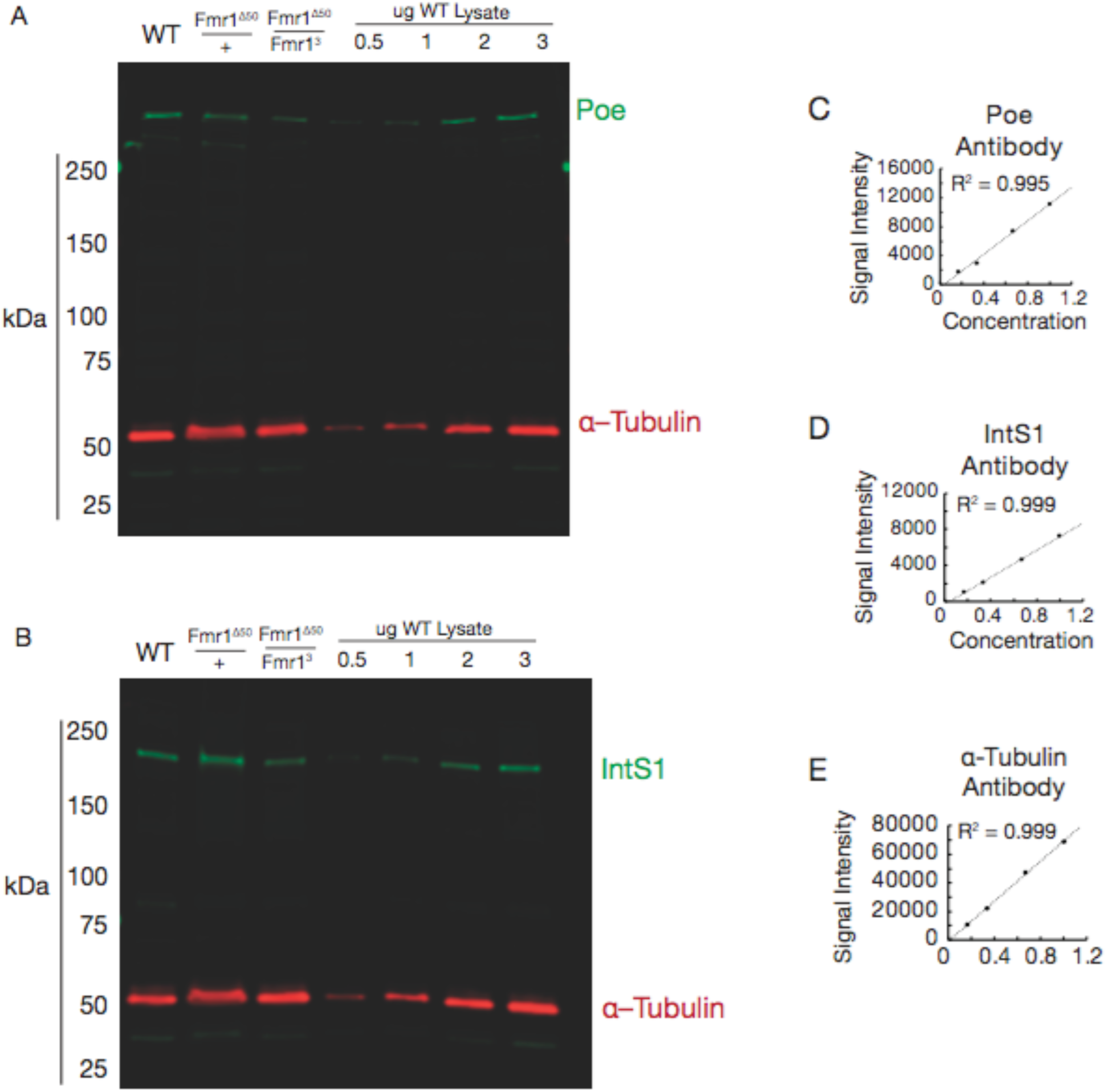
POE and INTS1 levels are reduced in Fmr1 mutant oocytes. **(A, B)** Full scans of Western blots of isolated unstored mature follicles of indicated control and Fmr1 mutant genotypes shown in Fig. 3E, F for **(A)** Poe and **(B)** IntS1 with α-tubulin serving as a loading control. **(C-E)** Linearity of Western blot signal using antibodies against **(C)** Poe, **(D)** IntS1, and **(E)** α-tubulin, as quantified using Image Studio software (LI-COR).

**Supplementary Figure 6.**
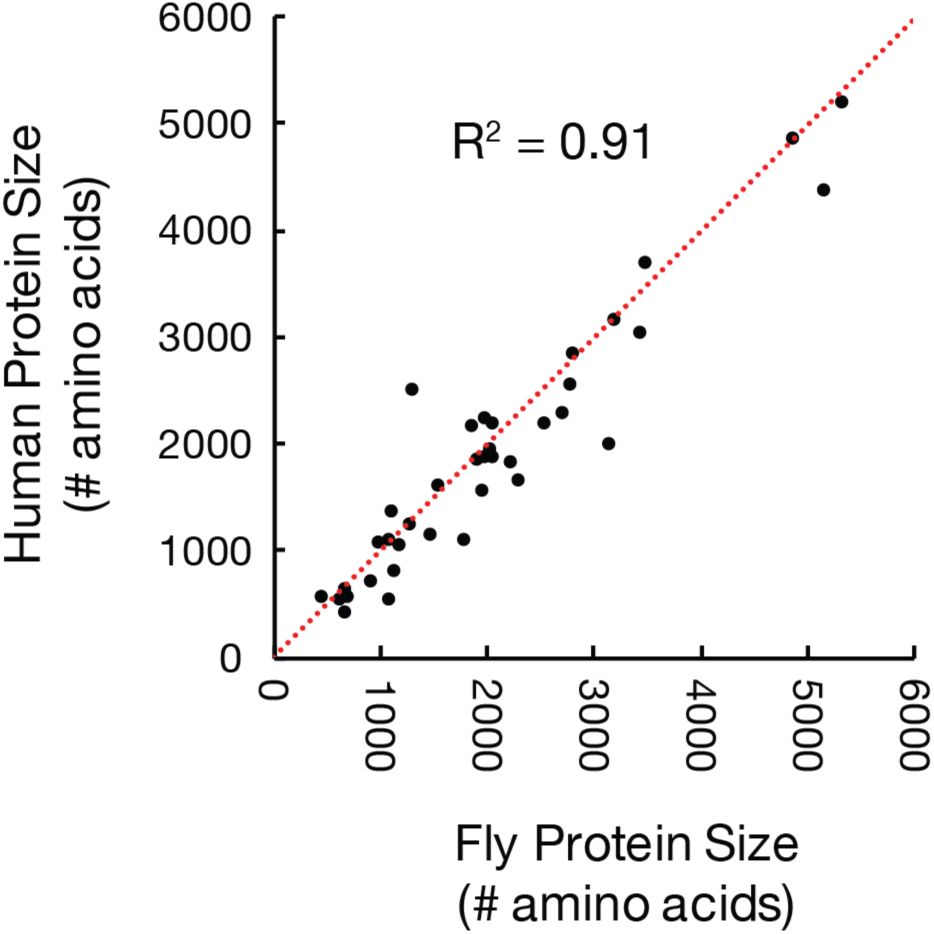
Conservation of size between *Drosophila* and humans among autism-related Fmr1 target proteins. Sizes of proteins encoded by Fmr1 target transcripts are well-conserved between *Drosophila* targets and homologous human autism-related genes (as described in Supplementary Table 3).

**Supplemental Table S1**

List of genes translationally downregulated in Fmr1 knockdown stored oocytes.

**Supplemental Table S2**

List of genes translationally upregulated in Fmr1 knockdown stored oocytes.

**Supplemental Table S3**

List of Fmr1 target genes with human homologs implicated in autism/intellectual disability.

**Supplemental Table S4**

Compiled ribosome profiling and mRNA-seq data.

## REFERENCES

1. Y. Feng et al., J. Neurosci. 17, 1539–1547 (1997).

2. C. T. Ashley, K. D. Wilkinson, D. Reines, S. T. Warren, Science. 262, 563–566 (1993).

3. G. Stefani, C. E. Fraser, J. C. Darnell, R. B. Darnell, J. Neurosci. 24, 7272–7276 (2004).

4. V. Brown et al., Cell. 107, 477–487 (2001).

5. Y. Feng et al., Mol. Cell. 1, 109–118 (1997).

6. A. J. Verkerk et al., Cell. 65, 905–914 (1991).

7. L. Wan, T. C. Dockendorff, T. A. Jongens, G. Dreyfuss, Molecular and Cellular Biology. 20, 8536–8547 (2000).

8. Y. Q. Zhang et al., Cell. 107, 591–603 (2001).

9. Cell. 78, 23–33 (1994).

10. R. L. Coffee, C. R. Tessier, E. A. Woodruff, K. Broadie, Disease Models & Mechanisms. 3, 471–485 (2010).

11. G. Deshpande, G. Calhoun, P. Schedl, Genetics. 174, 1287–1298 (2006).

12. L. Man, J. Lekovich, Z. Rosenwaks, J. Gerhardt, Front Mol Neurosci. 10, 290 (2017).

13. S. A. Barbee et al., Neuron. 52, 997–1009 (2006).

14. J. Dubnau et al., Curr. Biol. 13, 286–296 (2003).

15. A. Nakamura, R. Amikura, K. Hanyu, S. Kobayashi, Development. 128, 3233–3242 (2001).

16. M. Köhrmann et al., Mol. Biol. Cell. 10, 2945–2953 (1999).

17. M. R. Akins et al., Hum. Mol. Genet. 26, 192–209 (2017).

18. L. N. Antar, R. Afroz, J. B. Dictenberg, R. C. Carroll, G. J. Bassell, J. Neurosci. 24, 2648–2655 (2004).

19. A. Costa et al., Developmental Cell. 8, 331–342 (2005).

20. R. Rosario et al., PLoS ONE. 11, e0163987 (2016).

21. J. C. Darnell et al., Cell. 146, 247–261 (2011).

22. M. Ascano et al., Nature. 492, 382–386 (2012).

23. R. Tabet et al., Proc. Natl. Acad. Sci. U.S.A. 113, E3619–28 (2016).

24. L. Yang et al., Hum. Mol. Genet. 16, 1814–1820 (2007).

25. J. G. Dunn, C. K. Foo, N. G. Belletier, E. R. Gavis, J. S. Weissman, Elife. 2, e01179 (2013).

26. M. R. Wallace et al., Science. 249, 181–186 (1990).

27. S. M. Morris et al., JAMA Psychiatry. 73, 1276–1284 (2016).

28. J. Zhang et al., Hum. Genet. 136, 377–386 (2017).

29. S. H. Lelieveld et al., Nat. Neurosci. 19, 1194–1196 (2016).

30. D. Monies et al., Hum. Genet. 136, 921–939 (2017).

31. S. Moortgat et al., Eur. J. Hum. Genet. 121, 1085 (2017).

32. M. Bosshard et al., Sci Rep. 7, 15050 (2017).

33. C. M. Durand et al., Nat. Genet. 39, 25–27 (2007).

34. C. Sala, C. Vicidomini, I. Bigi, A. Mossa, C. Verpelli, J. Neurochem. 135, 849–858 (2015).

35. W. Pereanu et al., Nucleic Acids Res. 46, D1049–D1054 (2017).

36. S. Richards, T. Hillman, M. Stern, Genetics. 142, 1215–1223 (1996).

37. J. J. Fabrizio, G. Hime, S. K. Lemmon, C. Bazinet, Development. 125, 1833–1843 (1998).

38. B. Laggerbauer, D. Ostareck, E. M. Keidel, A. Ostareck-Lederer, U. Fischer, Hum. Mol. Genet. 10, 329–338 (2001).

39. Z. Li et al., Nucleic Acids Res. 29, 2276–2283 (2001).

40. E. G. Bechara et al., PLoS Biol. 7, e16 (2009).

41. M. Fähling et al., J. Biol. Chem. 284, 4255–4266 (2009).

42. A. Varshavsky, Protein Sci. 20, 1298–1345 (2011).

43. A. Khong et al., Mol. Cell. 68, 808–820.e5 (2017).

44. P. S. Estes, M. O’Shea, S. Clasen, D. C. Zarnescu, Mol. Cell. Neurosci. 39, 170–179 (2008).

45. D.-I. Kao, G. M. Aldridge, I. J. Weiler, W. T. Greenough, Proc. Natl. Acad. Sci. U.S.A. 107, 15601–15606 (2010).

46. R. Hagerman, P. Hagerman, Lancet Neurol. 12, 786–798 (2013).

47. S. D. Sullivan, C. Welt, S. Sherman, Semin. Reprod. Med. 29, 299–307 (2011).

## SUPPLEMENTAL REFERENCES

1. Ribosome profiling reveals pervasive and regulated stop codon readthrough in Drosophila melanogaster. 2, e01179 (2013).

2. Differential gene and transcript expression analysis of RNA-seq experiments with TopHat and Cufflinks. 7, 562–578 (2012).

3. Plastid: nucleotide-resolution analysis of next-generation sequencing and genomics data. 17, 958 (2016).

4. The developmental transcriptome of Drosophila melanogaster. 471, 473–479 (2011).

5. Drosophila Lk6 kinase controls phosphorylation of eukaryotic translation initiation factor 4E and promotes normal growth and development. 15, 19–23 (2005).

